# Using a fragment-based approach to identify novel chemical scaffolds targeting the dihydrofolate reductase (DHFR) from *Mycobacterium tuberculosis*

**DOI:** 10.1101/2020.03.30.016204

**Authors:** João Augusto Ribeiro, Alexander Hammer, Gerardo Andrés Libreros Zúñiga, Sair Maximo Chavez-Pacheco, Petros Tyrakis, Gabriel Stephani de Oliveira, Timothy Kirkman, Jamal El Bakali, Silvana Aparecida Rocco, Mauricio Luís Sforça, Roberto Parise-Filho, Anthony G. Coyne, Tom L Blundell, Chris Abell, Marcio Vinicius Bertacine Dias

## Abstract

Dihydrofolate reductase (DHFR), a key enzyme involved in folate metabolism, is a widely explored target in the treatment of cancer, immune diseases, bacteria and protozoa infections. Although several antifolates have proved successful in the treatment of infectious diseases, none have been developed to combat tuberculosis, despite the essentiality of *M. tuberculosis* DHFR (MtDHFR). Herein, we describe an integrated fragment-based drug discovery approach to target MtDHFR that has identified hits with scaffolds not yet explored in any previous drug design campaign for this enzyme. The application of a SAR by catalog strategy of an in house library for one of the identified fragments has led to a series of molecules that bind MtDHFR with low micromolar affinities. Crystal structures of MtDHFR in complex with compounds of this series demonstrated a novel binding mode that differs from other DHFR antifolates, thus opening perspectives for the development of novel and relevant MtDHFR inhibitors.

## Introduction

Dihydrofolate reductase (DHFR) is a key enzyme involved in the biosynthesis of tetrahydrofolate, an essential co-factor for the biosynthesis of purine and thymidine nucleotides and several amino acids^1^. DHFR is a well established target for human diseases, including cancer^2^, arthritis and other autoimmune diseases^3^, as well as for bacterial^4^ and protozoa infectious diseases, such as cystitis and malaria^5,6,7^. However, drugs such as antifolates used for human and infectious disease treatment differ considerably, due to differences in binding sites of microorganisms and human DHFRs; therefore the antifolate drugs targeting microorganisms generally have low side effects.

Despite the importance of DHFR inhibitors for human health, there is no drug developed which targets DHFR from *M. tuberculosis* (MtDHFR), possibly due to the low affinity of antifolates to this protein or the impermeability of *M. tuberculosis* cell wall to these compounds. However, several genes of the folate pathway in *M. tuberculosis* have been demonstrated to be essential. These include *folA*, which produces DHFR^8,9^ and for which there is extensive literature and using different strategies that have led to a large number of potential inhibitors^10,11,12^. Although methotrexate (MTX) is not used therapeutically against infections, it does inhibit *M. tuberculosis* growth *in vitro*^13^. *p*-Aminosalicylic acid (PAS), a second-line drug used in the treatment of tuberculosis, has been proposed to be an alternative substrate of the folate pathway and the product of the metabolism could also inhibit MtDHFR or the thymidylate synthase^14^.

Tuberculosis is one of the major infectious diseases and the cause of more deaths than any other bacterial species^15^. This disease has a high prevalence in a number of countries in Asia and Africa however this is now becoming a worldwide threat^16^. Various efforts have been focused on new strategies to combat tuberculosis, but only recently have two new drugs with novel action mechanisms been approved for use^17^. The discovery of new drugs against tuberculosis is particularly demanding because of the long regime therapy and the emergence of resistant strains against all used drugs. In this context, the development of new strategies to fight tuberculosis is an urgent need.

Fragment-based drug discovery (FBDD) is a successful approach that was introduced in companies such as Astex Pharmaceuticals, Abbott Laboratories and several other companies ^18^. Although this technique is still in development, about over 40 new drugs derived from fragments have been entered in clinical trials and four of them have been approved^19^. As the FBDD approach relies on the use of a small library with generally 1000 compounds, this technique has been widely applied in both the pharmaceutical industry^20^ and in academia ^21,22^. The FBDD approach involves the identification of low molecular weight compounds, with about 200-250 Da through a range of biophysical^23^, biochemical^24^ and crystallographic methods^25^. Once the binding of the compounds (fragments) to the target has been validated and the binding mode elucidated, these small compounds are a very attractive starting point for the synthesis of high-affinity compounds^26^. Fragments identified from a fragment screen generally have affinities between 2 mM and 100 µM, and these can be quickly elaborated by organic synthesis to optimise interactions and give lead compounds that have sub-micromolar affinities for the target enzyme^26^. Although FBDD has been applied to several targets from *M. tuberculosis*^27,28,29,30^ and new compounds have been identified to inhibit the growth of this bacterium^29,31^, this technique has not been applied to *Mt*DHFR or any other DHFR.

In this work, the FBDD approach has been successfully applied using a range biophysical techniques, including Differential Scanning Fluorimetry (DSF), Isothermal Titration Calorimetry (ITC), ligand-based Nuclear Magnetic Resonance (NMR) and Crystallography to identify compounds that have completely novel scaffolds that bind to the active site of the *Mt*DHFR with 2 mM to 80 µM affinities. In addition, one of these compounds was used as a starting point for further elaboration where from an in house library of compounds low micromolar affinity compounds with a completely novel scaffold for a DHFR inhibitors were identified. These compounds that target DHFR do not contain the 2,4-diaminopyrimidine moiety which is found widely in DHFR inhibitors. Furthermore, 13 X-ray crystal structures of MtDHFR in complex with different molecules were solved and these will prove to be crucial in the design of new antifolates that could contribute to further tuberculosis drug discovery campaigns.

## Material and Methods

### Protein production and preparation

MtDHFR was produced according to the protocol established by Dias et al.^32^ and that of Ribeiro et al.^33^ with a few modifications. Briefly, the *folA* gene from *M. tuberculosis* inserted into pET28(b) expression vector was transformed in BL21(DE3) cells and the expression was induced using 0.2 mM IPTG at 18 °C overnight by constant agitation at 220 rpm. The cells were resuspended in 20 mM potassium phosphate buffer, pH 7.5, 50 mM KCl (Buffer A) and disrupted by sonification. The cell debris was removed by centrifugation at 14,000 rpm and the supernatant was loaded onto an IMAC column (GE Healthcare) charged with nickel. The protein was eluted using buffer A plus 500 mM of imidazole (buffer B), using a linear gradient of buffer B to increase the concentration of imidazole on an ÄKTA purifier system (GE Healthcare). The protein was concentrated and further purified using an S200 16/60 gel filtration chromatography column (GE Healthcare) previously equilibrated with buffer A. The protein was concentrated to at least 10 mg/mL^-1^ and stored at −80°C.

### Differential Scanning Fluorimetry (DSF)

DSF assays were performed in a PCR iCycler (BioRad) coupled to an iQ5 Multicolor Real-Time PCR Detection System (Bio-RAD). 96 well plates were used with each well containing a total volume of 100 µL consisting of 95% (v/v) buffer A, 5% (v/v) DMSO, 2.5x SYPRO Orange® dye (Invitrogen) and 1 mM NADPH (Sigma-Aldrich). The final concentration of MtDHFR in each well was 3 µM. 80 wells of the plate were used to screen different fragments, which had a concentration of 5 mM, 8 wells were used as positive controls (5 mM of trimethoprim) and 8 wells were used only in the presence of 5% DMSO as “negative control”. 1,250 fragment-like compounds from the Maybridge RO3 library were tested. For the assays, the plates were heated from 25°C to 75°C at increments of 0.5°C per minute. The fluorescence intensity of SYPRO Orange® dye was monitored, with excitation-emission wavelengths of 490/575 nm, as a function of temperature. The ΔTm was calculated from the difference of the average values of all “negative controls” (reference) and that of the protein in the presence of fragments. A positive hit was defined as a compound that increased the melting temperature by ≥ 0.5°C compared to that of the reference compound.

### STD-NMR Assays

For NMR experiments, the fragments were initially dissolved using DMSO-*d*_*6*_ and further diluted using D_2_O at a final concentration of 500 µM and the protein in buffer A was diluted to 5 µM. The NMR spectra were obtained using an Agilent DD2 500 MHz spectrometer equipped with a triple-resonance probe at 298 K. The 1D STD-NMR spectra were obtained by the subtraction of saturated spectra (on resonance) from the reference spectra (off-resonance) after identical processing and phasing. The subtraction of on-resonance spectra from off-resonance spectra was performed automatically by phase cycling using the dpfgse_satzfer pulse sequence implemented in the VNMRJ software (Agilent). The 1D STD-NMR spectrum was acquired using 8192 scans with a selective irradiation frequency of protein at −0.5 ppm for on-resonance and at 30 ppm for off-resonance. 40 Gauss-shaped pulses of 50 ms separated by a 1 ms of delay were applied to the protein. The total length of the saturation train was 2.04 s. A T_2_ filter was applied to eliminating all protein background. The off-resonance spectrum was used as a reference spectrum and acquired with 4096 scans keeping all the other parameters equal to the 1D STD-NMR spectrum. For the group epitope mapping analysis, the STD enhancements were determined by the integrals of individual protons of the ligands in the 1D STD_NMR spectrum divided by the integral of the same signals at the reference spectrum. The strongest STD enhancements in each spectrum are normalized and assigned to 100%.

### Isothermal Titration Calorimetry (ITC)

ITC assays were performed on a MicroCal VP-ITC 2000 or iTC200 (Malvern). For both, cell and syringe solutions contained 95% (v/v) buffer A, 1 mM NADPH and 5% (v/v) DMSO. The cell solution contained between 30 and 50 µM of MtDHFR for experiments performed using VP-ITC 2000 or between 100 to 160 µM for experiments using an iTC200. All tested compounds were dissolved at a concentration ranging from 2 to 10 mM in buffer A in the presence of in 5% of DMSO and 1 mM of NADPH to avoid the buffer mismatch. All solutions were degassed prior to the experiments and the temperature used in the cell was 26 °C. The titration constituted of 20 to 25 injections using different volumes for VP-ITC (first injection of 3 µL followed by 24 injections of 10 µL) and iTC200 (first injection of 0.2 µL followed by 19 injections of 2.2 µL). The heats of dilutions were subtracted from the titration data prior to curve fitting. Most of the obtained curves were fitted by nonlinear least-squares regression using the non-interacting one-site model using Origin 7.0 (Microcal).

### Crystallography and structure analysis

The crystallisation of MtDHFR was performed by the vapor diffusion method using a hanging-drop strategy and the protocol described by Ribeiro et al.^33^ Briefly, 1 µL of MtDHFR at 10 mg/mL, previously incubated with 10 mM of NADPH, was mixed in a coverslip with 1µL of a crystallisation condition constituted by 1.6 M ammonium sulfate, 100 mM MES (2-(N-morpholino) ethanesulfonic acid), pH 6.5, 10 mM CoCl_2_ and inverted down against the 300 µL of crystallisation solution added in the wells of Linbro plate (Hampton Research). Bipyramidal crystals of MtDHFR appeared after 3-4 days at a temperature of 18 °C. Complexes of the holoenzyme (MtDHFR:NADPH) with fragments and compounds were obtained by co-crystallisation or soaking as described in Table 1. For co-crystallisation, MtDHFR was prior incubated for about 30 minutes on ice in the presence of 20 to 60 mM of different fragments. For soaking experiments, about 50 to 100 mM of a compound solution was added to a drop containing crystals of holoenzyme for 10 minutes to 20 hours, prior to crystal harvesting and freezing, depending on the damage to the crystal caused by the compound.

**Table 1.**
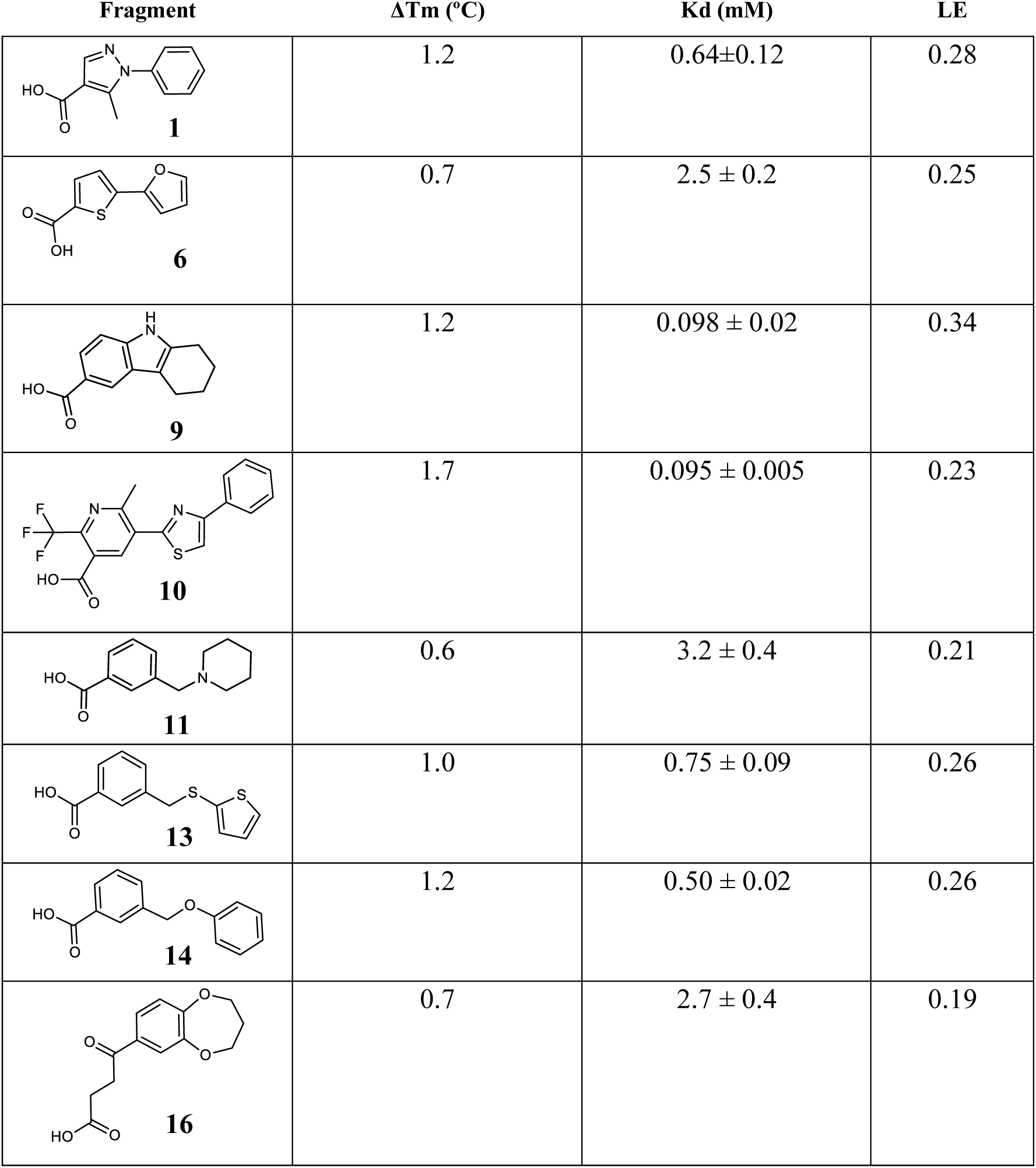
Calorimetric analysis of fragments against MtDHFR

| Fragment | $\Delta T_m$ (°C) | $K_d$ (mM) | LE |
| --- | --- | --- | --- |
| **1** | 1.2 | 0.64±0.12 | 0.28 |
| **6** | 0.7 | 2.5 ± 0.2 | 0.25 |
| **9** | 1.2 | 0.098 ± 0.02 | 0.34 |
| **10** | 1.7 | 0.095 ± 0.005 | 0.23 |
| **11** | 0.6 | 3.2 ± 0.4 | 0.21 |
| **13** | 1.0 | 0.75 ± 0.09 | 0.26 |
| **14** | 1.2 | 0.50 ± 0.02 | 0.26 |
| **16** | 0.7 | 2.7 ± 0.4 | 0.19 |

X-ray data collection of MtDHFR in complex with different fragments and compounds was achieved using different synchrotron sources, including the Laboratório Nacional de Luz Síncrotron (Brazil) and Diamond as described in Table 1. The data processing was performed using XDS^34^ and scaled by Aimless^35^ from the CCP4 suite^36^. The structures were solved by molecular replacement through the program using Phaser MR^37^ using the PDB entry 1DF7^38^ included in the Phenix crystallographic suite^39^. The crystallographic refinement was performed by the program Phenix.Refine^40^ followed by manual inspection using COOT^41^. The stereochemical quality of the models was checked using the program MolProbilty^42^ and the figures were prepared using PyMol (The PyMOL Molecular Graphics System, Version 2.0 Schrödinger, LLC).

## Results and discussion

### Screening by DSF and validation by 1D STD-NMR

MtDHFR was screened against a library of 1250 compounds selected following the rule of three^43^ using a fluorescence-based thermal shift method in the presence of 1 mM of NADPH. Trimethoprim (TMP), which gave a ΔTm of +10°C (Supplementary Figure 1) was used as a positive control. However, early in the screening process, it was observed that T_m_ of negative controls varied slightly from well to well making it difficult to determine the threshold for selecting fragments. Hence, to define a threshold, 105 observations of T_m_ of MtDHFR:NADPH in 5% of DMSO were collected as “negative controls” and in order to verify whether these data were describing a normal distribution, two goodness-of-fit tests (Kolmogorov-Smirnov and Chi-Squared tests) were performed. Both of the tests suggested that these data are normally distributed, with a mean T_m_ of 53.2 °C and standard deviation (σ) of 0.7 °C, up to a significance level of α=0.02. Using this information, a 99% confidence interval was constructed, giving a range of temperatures between 53.0 and 53.3 °C, within which 99% of the mean T_m_ values of the negative controls should lie. From this analysis, those fragments that produced flat, markedly-irregular melting curves were excluded since the ΔT_m_ could not be calculated. Positive hits were then defined as fragments that gave a ΔTm of ≥ 0.5°C above 53.4 °C. About 37 statistically positive hits (Supplementary figure 2), which represent about 2.8% of the library with significant chemical diversity, were identified when screened at a fragment concentration of 10 mM. This hit rate is similar to that obtained in several other FBDD campaigns^27,44,24^. The remaining molecules of the library did not significantly stabilise the MtDHFR or displayed a negative ΔT_m_, indicating a stabilisation of the unfolded state of MtDHFR, consequently, these molecules were also not considered for further analysis.

A large number of identified compounds shared similar features, including a carboxylic acid at the *meta* position of a benzene ring, which is generally linked by a single bond to another comprising two heterocyclic rings (Figure 1). Interestingly, it has been reported that compounds with stilbenoid, deoxybenzoin and chalcone moieties, which have similar scaffolds to some of our compounds, also showed affinity to *E. coli* DHFR and had an uncompetitive mechanism of inhibition^45^ (Supplementary figure 2).

**Figure 1.**
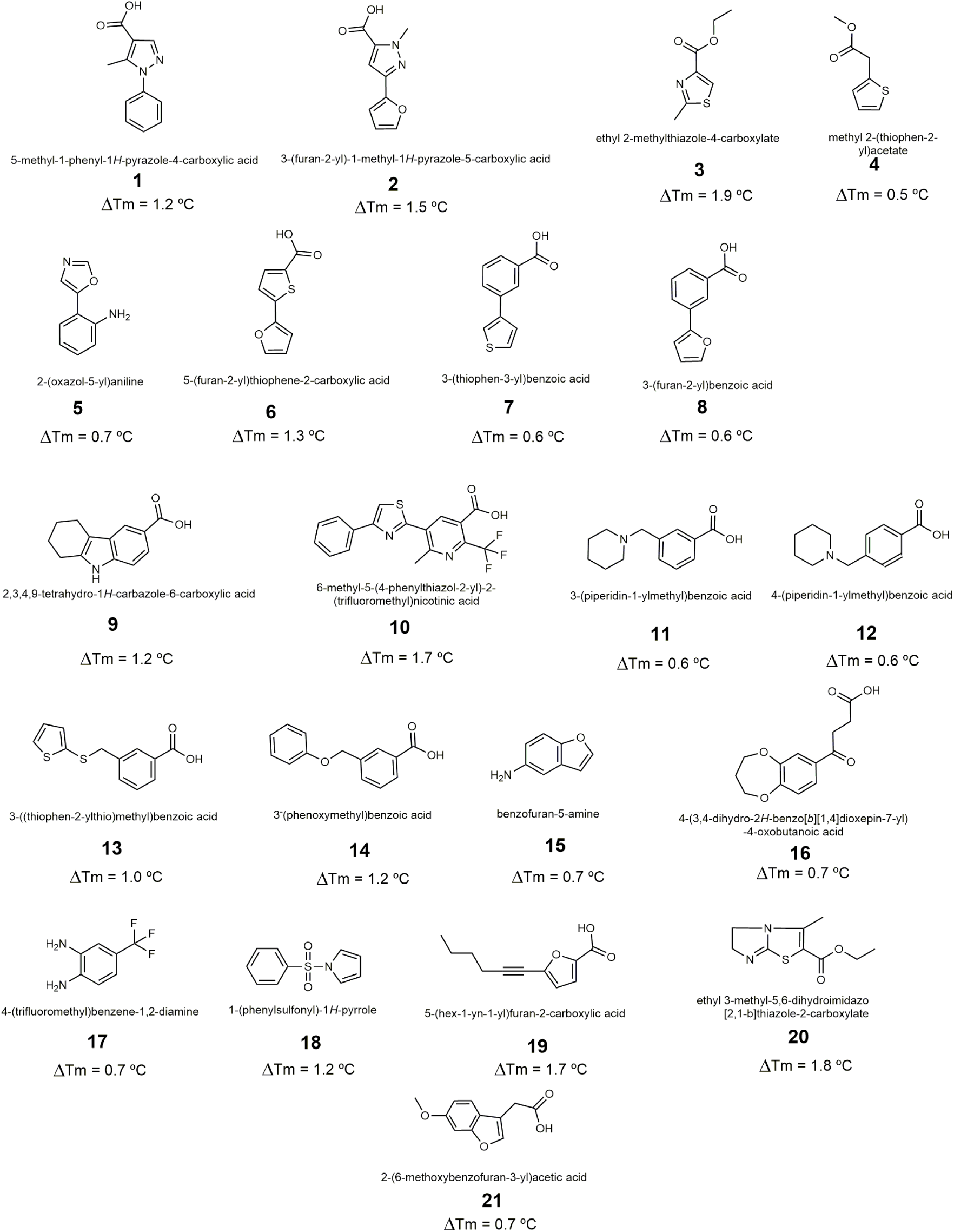
Fragments validated by STD-NMR. The ΔTm is included below each structure

From 37 molecules that stabilised MtDHFR above than 0.5 °C in the DSF assay, we rescreened 30 fragments (Supplementary Figure 2) using 1D STD-NMR and confirmed 21 fragments molecules (Supplementary Figure 3) that interact with the protein, which represents about 70% of the molecules (Figure 1) indicating a strong accuracy of statistic method applied for the selection of the fragments.

Although 1D STD-NMR is routinely used to rescreen the hits identified by DSF, we also used this technique to identify epitopes from the fragments that are essential for the interaction with the target^46^. The identification of epitopes is possible because of the proximity of the functional groups of the side chain of amino acid residues of the target, and the degree to which this occurs is dependent on the time the ligand spends in contact with the protein.^47^ 1D NMR spectra of solutions containing the fragments and the holoenzyme were measured to discriminate the signals. The fragment interactions with the target were confirmed by the proton signals in the difference spectra (on-resonance – off-resonance). The data of difference were normalised and the signal/noise rate was calculate and a threshold of ≥ 5 for this relation. In addition, we have attributed to the signal of proton as the more intense for STD enhancements and consequently, we considered it as 100% and they have been used as a reference for the calculation of the intensity percentage for the other protons of the ligand. Proton signals were satisfactorily confirmed for the 21 fragments and the corresponding protons attributed in the spectra are identified by numbers in the structure. The remaining signals in the reference spectra belongs mainly to the coenzyme NADPH (Supplementary Figure 3). An interesting observation obtained by the 1D STD-NMR signal was that the benzoic acid for most of the fragments should have close contacts with the protein since the STD enhancement for this group is nearly 100 %. Consequently, this region should bind tightly to the target and have a key role in the interaction. In addition, it is observed that most of the compounds that have this benzoic acid moiety are linked to other heterocycles, for example, thiophene (**7** and **13**), furan (**8**), carbazole (**9**) and non-aromatic heterocycles, such as piperidine (**11** and **12**). The fragments **11** and **12** are similar molecules and they differ only in the position of the carboxylic acid at the benzene ring, which for **11** is at *meta* position and in **13** is *para*. As observed for most of the fragments with benzoic acid, **11** has an intense signal of protons and consequently, the *meta* configuration of the carboxylic acid attached to the benzene ring could be a key factor for the strong interaction. The crucial role of the benzoic acid for the interaction suggested that this group could perform an ionic interaction with one of the arginines within the active site of MtDHFR, such as Arg60, mimicking the interaction of glutamate moieties of dihydrofolate. The *meta* position of the carboxylic acid could contribute to orientate this group towards the positive charges of the active-site cavity. In addition, the importance of the carboxylic acid attached to a ring could also be observed for compounds **1, 2, 6** and **10**, where these instead of having benzene ring, have the carboxylic acid attached to a heterocycle such as a pyrazole, thiophene, and pyridine so that a maximum resonance could be observed for carbons of the ring. In addition, the 1D STD-NMR spectra also suggest the importance of different rings linked since the transference of resonance is highly significant. Moreover, there is a clear preference of aromatic rings, including both benzene or heterocycles, including pyrazole (**1**), thiazole (**3**), oxazole (**5**) and pyrrole (**18**). Interestingly, compound **16**, which has the benzoic acid fused with a dioxepine, but the carboxylic acid attached to the benzene ring through a 4-oxo-butyl linker could have a key role in the interaction with the protein.

In addition to the fragments identified above, a number of other structures were confirmed to interact with MtDHFR. For example, another group of fragments comprises of small compounds with neither linked rings nor carboxylic acid (**3, 4, 15, 17**). Although two fragments (**5** and **18**) have linked rings, they do not have carboxylic acid groups.

### Calorimetric analysis of fragments selected as hits for DHFR

The 21 fragments that were confirmed to bind to MtDHFR through DSF and 1D STD-NMR, were further characterised by ITC (supplementary table 1 and supplementary Figure 4). For 5 compounds, the thermodynamic parameters were not determined due to their low solubility or there being not enough heat released in the ITC experiment conditions. For 8 molecules the affinity was too low to estimate the LE (ligand efficiency) or had a too-high entropic penalty, which strongly affects the ΔG (Supplementary Table 1). Table 1 shows the affinity and LE for the 8 fragments. In addition, except for the titration with **1**, all thermograms were fitted using a unique site model, indicating that the MtDHFR active site is occupied only by one ligand molecule (Supplementary figure 4). Interestingly, the thermogram curve for **1** indicated a sequential site model, suggesting that this molecule might bind in two distinct regions of the active site. **9** and **10** were the fragments that have the highest affinity among the tested molecules, with 95 and 98 µM, respectively, although **9** also has the highest LE, 0.34.

### Structural basis of the interaction of MtDHFR with selected fragments

Once the fragments that interact with MtDHFR had been characterised biophysically, the next step was to define the structure of the fragments with MtDHFR in order to examine their interactions with the proteins. A total of 9 crystal structures were obtained with a resolution ranging from 2.5 to 1.7Å for MtDHFR in complex with different fragments (Supplementary Table 2). Although the binding affinity for fragments **3** and **16** were not able to be measured, their interaction modes with MtDHFR, despite the low occupancy of these molecules was obtained. The structures obtained for most complexes of MtDHFR:NADPH with fragments was through co-crystallisation, in which the formation of the crystal was not inhibited by the high concentration of compounds (about 15 to 40 mM). The complex of MtDHFR:NADPH and **10** was obtained by soaking in the crystallisation condition described by Dias et al., 2014^32^. The crystals of MtDHFR:NADPH in complex with different fragments, except the complex of MtDHFR:NADPH with **10**, belong to space group P2_1_2_1_2_1_ with 2 molecules in the asymmetric unit. For most of the complexes, we observed both protomer active sites occupied. Supplementary Table 2 shows the data processing, refinement and quality of the stereochemistry of the structures of the complex with fragments. All structures of MtDHFR:NADPH in complex with fragments have a closed conformation with the NADPH engaged in the active site as described previously^32,38^. In addition, it was observed that the protein did not undergo significant conformational changes, even in the residues of the dihydrofolate binding pocket (Supplementary Figure 5). To facilitate the description of fragment interaction with the protein, we distinguished four different sub-regions of the active site groove of MtDHFR:NADPH (Figure 2): (1) active site entrance, with a positive charge, (2) central region of the active site or PABA binding site, a slightly apolar region, (3) dipyrimidine binding site, with strong negative charges; and (4) the glycerol binding site, a specific region found in MtDHFR, but not observed in human DHFR (hDHFR)^38^.

**Figure 2.**
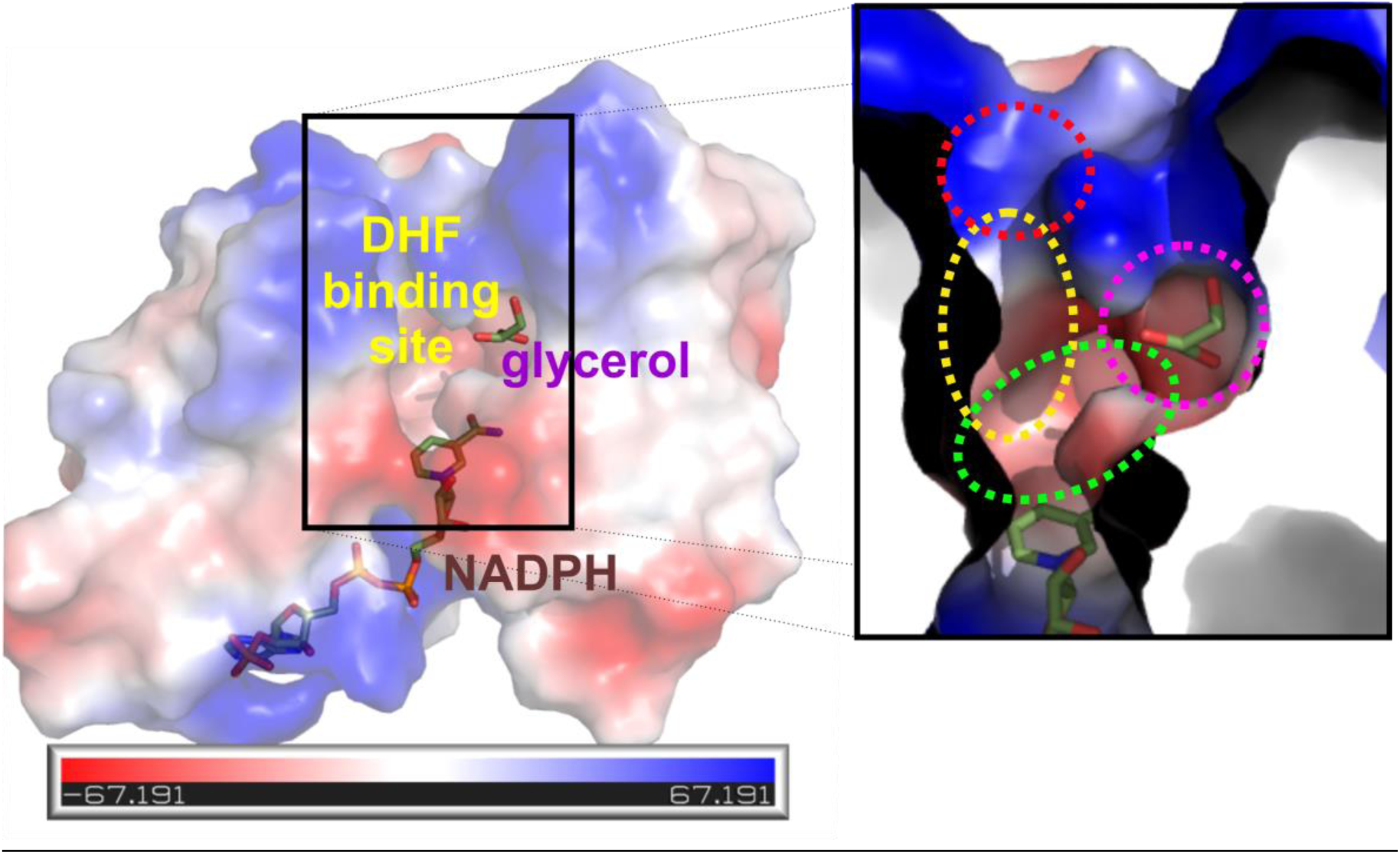
Active site groove of MtDHFR and subregions. The figure shows the DHF and glycerol binding cavities. In the inset, we have specified 4 subregions of the active site. 1 (in red) is a positively charged region that binds the glutamate moiety of DHF; 2 (in yellow), the central region, where the PABA moiety sits, that is apolar, 3 (in green), where the positive dipyrimidine ring of DHF stacking with nicotinamide moiety of NADPH and finally 4 (in magenta), the glycerol binding site, an exclusive pocket of MtDHFR in comparison to human DHFR.

All the structures of MtDHFR:NADPH in complex with fragments show electron density for fragments in three regions of MtDHFR:NADPH, which includes the entrance (1), PABA binding site (2) and glycerol binding site (4); surprisingly we did not observe any fragment completely occupying the dipyrimidine binding site (3), which is one of the most explored regions of DHFR in the development of new inhibitors. Most of the compounds that have the carboxylic acid linked to a phenyl or a heterocyclic ring indicate strong interaction with the active site entrance (region 1), where the carboxylic acid is interacting through ionic interaction with Arg60. The interactions of the compounds in that region explain and corroborate our results from 1D STD-NMR, which indicates a 100 % proton transference of this region and the high enthalpic contribution for the binding, characterising a strong polar interaction. The importance of a second ring (phenyl or heterocyclic) could also be confirmed since in most cases these rings exhibit π-interactions with Phe31 on the PABA binding site (sub-region 2), as well as interactions with other residues, such as Leu50 and Ile94 (Figure 3). These interactions also confirm the results of ITC indicating high entropic contributions for the interaction of fragments (Supplementary Table 1). In addition, because of the proximity of the second ring of these compounds with the nicotinamide group of NADPH, we can also observe an edge-π interaction with this group and then slightly reaching the subregion 3.

**Figure 3.**
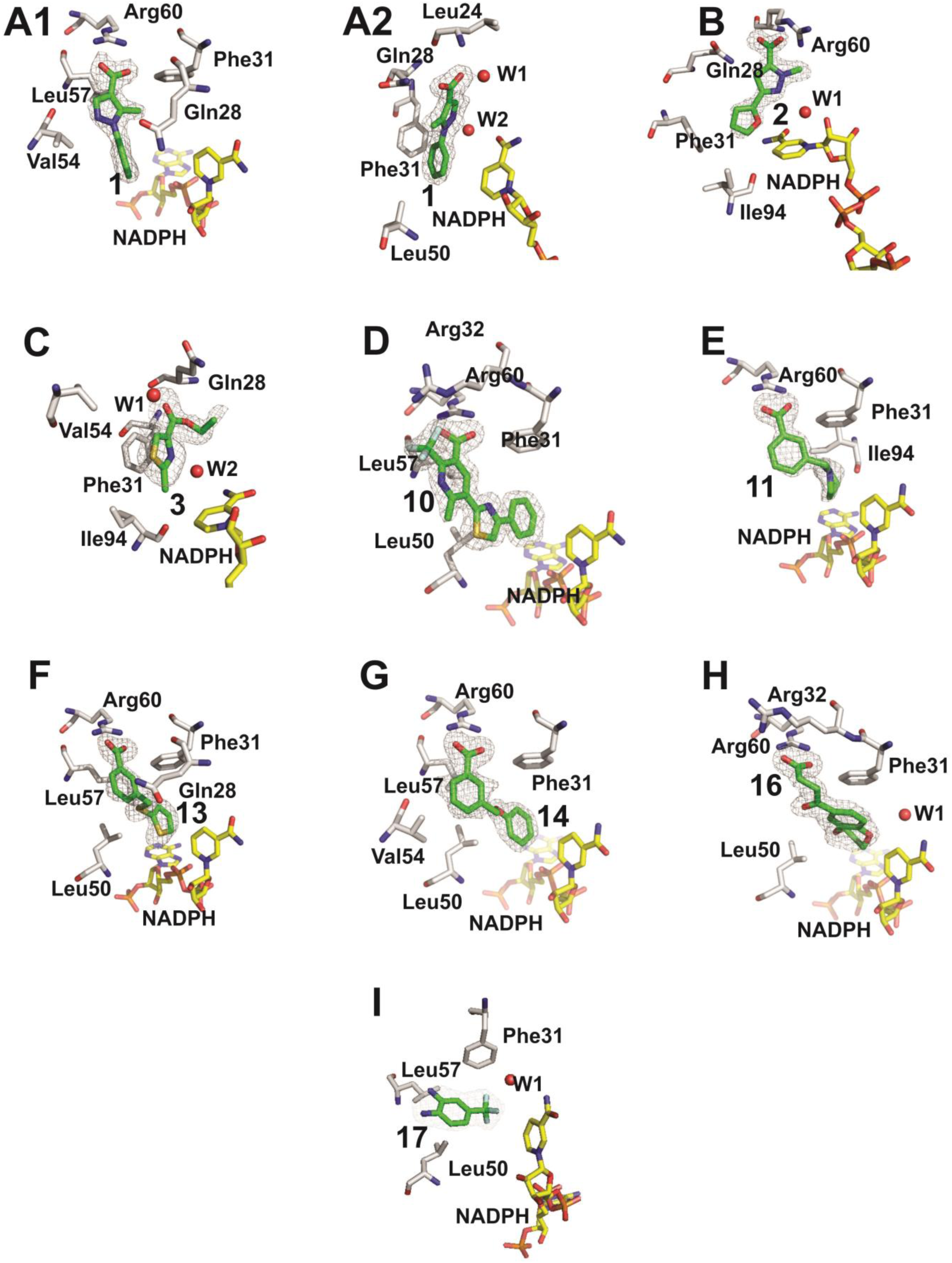
Crystal structures and binding modes of nine fragments that bind to MtDHFR:NADPH. (A) **1**. Fragment **1** binds in two different pockets of MtDHFR (**A1** and **A2**) (PDB entry 6VS5); (B) Fragment **2** (PDB entry 6VS6); (C) Fragment **3** (PDB entry 6VS8); (D) Fragment **10** (PDB entry 6VVB); (E) Fragment **11** (PDB entry 6VS9); (F) Fragment **13** (PDB entry 6VSD); (G) Fragment **14** (PDB entry 6VSE); (H) Fragment **16** (PDB entry 6VSF); (I) Fragment **17** (PDB entry 6VSG). The electron density contorns is from a 2fo-fc map with 1σ. W1 are water molecules on the active site cavity.

Fragment **2** was unique in that it did not interact with Arg60 but rather is hydrogen-bonded to the side chain of Gln28 even having the carboxylic acid in *meta* position while maintaining the interaction with Phe31 through its furan group as do the other fragments from this group. Interestingly, fragment **1** was unique in occupying the glycerol-binding site (4), while its carboxylic acid makes a hydrogen bond with the side chain of Gln28. Since the glycerol-binding site is a feature observed in MtDHFR in contrast to hDHFR, this fragment could be an excellent starting point to obtain specific drug leads against tuberculosis. As expected, an electron density for fragment **1** was also observed in the PABA binding site (2), corroborating the results of ITC that indicated that this molecule has a sequential binding mode. However, as the average B-factor for this second molecule is higher than the first one, probably the binding just occurs after the complete occupation of the first site.

Although there is a structural similarity of **6-8** to **1** and **2**, we were not able to obtain the structures of these fragments even though they have a carboxylic acid group. As the affinities of **2, 5-8** are low, we suggest that a higher affinity interaction of this fragment scaffold would be necessary for the association of “*meta*-position” in a five-membered ring with a phenyl ring, a characteristic observed only for **1**. Interestingly **1** and **2** are also very similar compounds, but the binding modes of these two molecules are quite different, indicating that the phenyl and furan groups, as well as the position of the methyl group attached to the pyrazole ring, might have a strong influence on the orientation of these compounds in the active site of the enzyme.

Fragments **3** and **17** were the only ligands in the series that do not have a carboxylic acid for which we were able to obtain the crystal structures; however, we could observe the binding mode in only one protomer of the asymmetric unit for both compounds and we were not able to obtain the affinity of these compounds by ITC. These two molecules have distinct structures since **3** is a thiazole ethyl ester and **17** is a trifluoromethylphenyl diamine. Interestingly, both molecules bind in the PABA binding site (3), despite their different orientations and make only non-polar contacts with the active site residues.

Fragment **10** is the largest for which we have obtained the crystal structure complex with MtDHFR:NADPH. Although its thiazole ring is binding in the same position of the PABA moiety of MTX, the phenyl ring reaches a deeper point in the active site of MtDHFR and sits close to the pyrimidine moiety of MTX stabilising several non-polar contacts with this active site region (Supplementary Figure 6).

The chemical structures of the fragments **13** and **14** are similar and they differ in a second ring in which **13** and **14** have a thiophenyl-thiomethyl and a phenoxymethyl and they bind in the similar position at the active site of MtDHFR (although the twist caused by the sulfur atom in contrast to the oxygen in the linker between the two rings). The similar pose of these fragments in the active site of MtDHFR was expected since their affinities (Kd) were also similar, 0.752 and 0.502 mM for **13** and **14**, respectively. Interestingly, although these compounds are smaller than **10**, the thiophene and the phenyl ring sits in the same position of the phenyl ring as compound **10** and the linker between the two rings adopts the position of the PABA moiety of MTX (Figure not shown).

The role of the carboxylic acid in the meta position could be further validated only by the structure of MtDHFR:NADPH in complex with **11.** Even though we performed several attempts to obtain the structure for the complex with **12**, crystals were not forthcoming. Although these two compounds have planar piperidine rings, **11** is able to perform the ionic interaction with Arg60 through its “*meta*” carboxylic acid. The variation in the position of the carboxylic acid attached to the phenyl rings causes a three-fold difference of affinity between the two compounds.

Finally, **16** is the only compound to have a benzodioxepinyl group and an oxobutanoic acid attached to position 7 of the benzyl group. Although the restraints caused by the fusion of the benzo and dioxepinyl rings on the flexibility of the oxobutanoic acid moiety, this fragment also interacts with the Arg60 through the carboxylic acid. However, the benzodioxepin moiety does not superpose well with the position of the other fragments with different linkers between the two rings and instead, only the dioxepinyl ring sits in the same binding site as the phenyl group of compound **10.**

### Applying SAR by catalog using Fragment 1 to obtain new leads with a low-micromolar affinity against MtDHFR

Fragment **1** binds to the glycerol pocket of MtDHFR with a K_D_ of 0.640 mM (LE = 0.28) and it was selected for further elaboration and optimisation. Since the aryl rings of the compound play an important role in the affinity to MtDHFR, it was decided that both rings should be maintained during any further optimisation. Based on fragment **1**, a campaign of SAR was explored using an in-house library (Supplementary methods). Initially, DSF was used to identify compounds that had a higher ΔT_m_ than fragment **1** and a number of compounds were identified and the affinities of these were assessed using ITC (Table 2 and Supplementary Figure 7). These compounds have the scaffold of Fragment 1 and an indole moiety connected with different linkers (Table 2). In addition, the crystal structures of MtDHFR:NADPH in complex with four of these compounds were obtained (Figure 4 and Supplementary Table 3). Interestingly the ITC indicated a significant increase of affinity for three molecules, in the range of 10, 20, and 40-fold greater than the initial fragment and the LE decreased only slightly for the compounds **1c** and **1d**, respectively (Table 2). The best of these compounds (**1d**) had a K_D_ of 17 µM and has a propylamine linker. The complexes of MtDHFR:NADPH with these 4 compounds had their crystallographic structures solved and the gain of the affinity of **1d** was provided by the oxygen of the propanamide group linker that is able to form a new hydrogen bonding interaction with Gln28 side chain in contrast to the other compounds of this series (Figure 4). The analysis of the structures of the complexes indicates that probably a shorter linker is favorable to accommodating these compounds in the active site and performing π-interaction with Phe31. However, because of its flexibility, the linker can adopt different conformations as well as the indole ring, although this always performs a π interaction with a Phe31. In contrast, the fluorine atom attached to the aryl ring of the compound **1a** seems to strongly influence the affinity of this compound as it has the lowerest affinity amongst these series. The addition of the methyl group on the indole ring of compound **1c** also could decrease the affinity of this compound in contrast to the **1d**. The thermodynamic analysis of **1d** indicates a high entropic penalty in comparison with the other compounds of the series (Supplementary Figure 7).

**Table 2.**
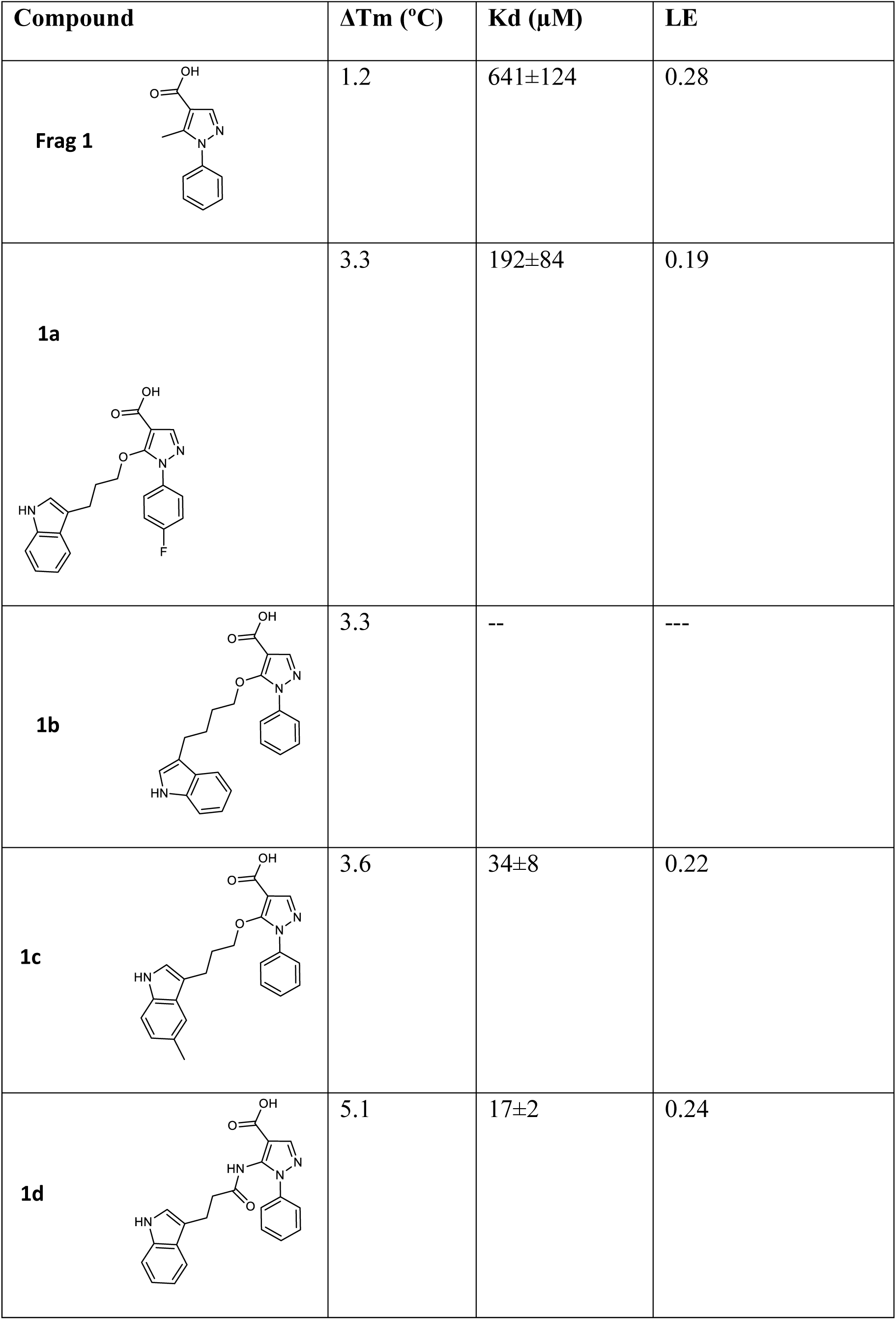
Calorimetric analysis of derivatives from Fragment **1**.

| Compound | $\Delta T_m$ (°C) | K<sub>d</sub> (μM) | LE |
| --- | --- | --- | --- |
| **Frag 1** | 1.2 | 641±124 | 0.28 |
| **1a** | 3.3 | 192±84 | 0.19 |
| **1b** | 3.3 | -- | --- |
| **1c** | 3.6 | 34±8 | 0.22 |
| **1d** | 5.1 | 17±2 | 0.24 |

**Figure 4.**
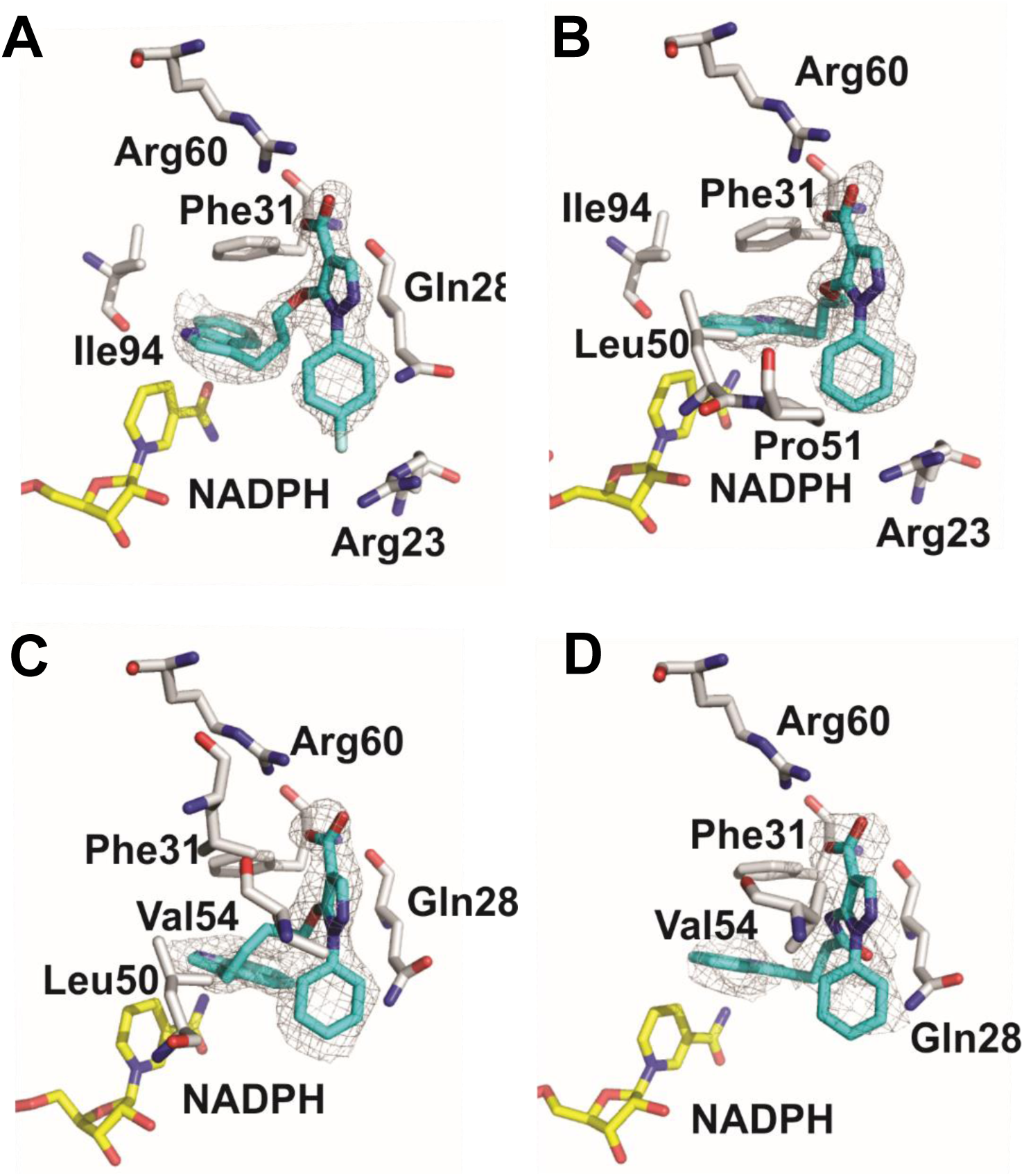
Crystallographic structures of compounds derived from Frag. 1. A) **1a** (PDB entry 6VV6; B) **1b** (PDB entry 6VV9); C) **1c** (PDB entry 6VV7) D) **1d** (PDB entry 6VV8). The electron density is based on a 2Fo-Fc map with a 1σ contours.

However, these four compounds have lost their interaction with the glycerol binding pocket adopting the binding mode of position 1, which it is shown in Figure 3A1, most likely due to the strong interaction between the indole group and Phe31, and the favourable interaction of the carboxylic acid with the Arg60 at the active site entrance. Further optimisation is necessary to maintain chemical functional groups in this pocket.

The compounds based on fragment **1** show that the development of new DHFR inhibitors based on a pyrimidine free scaffold is feasible and a more stepwise approach for fragment growing, expanding into the rest of the active site whilst maintaining the glycerol binding site interaction should be possible. As the glycerol binding site is a specific feature of MtDHFR, selective compounds targeting *M. tuberculosis* rather than the human DHFR could be obtained.

## Conclusions

We have successfully applied an FBDD approach to the enzyme MtDHFR and have identified and confirmed the interactions of 21 molecules. This is the first report of an FBDD campaign against any DHFR and surprisingly, many of the identified compounds have completely novel scaffolds not reported for any DHFR. Through subsequent screening of an in-house library, derivatives of one of the molecules (**1**) were identified and binding assessed, showing improved affinities from 600 µM to 17 µM, although interactions with the glycerol binding site were lost. However, the identification of these lead molecules opens possibilities in the identification of a completely novel series of compounds targeting MtDHFR and that could move forward to *in vivo* studies.

## Acknowledgments

This research was supported by FAPESP (Sao Paulo Research Foundation, grants 2010/15971-3, 2014/09188-8 and 2018/00351-1, The Bill & Melinda Gates Foundation (Grant RG48788 MAAG/555 and PHZF/121), CNPq grant 442021/2014-3 and Royal Society of Chemistry grant RF-17-9399. JAR and SMCP received fellowships from the FAPESP, fellowship 13/15906-5 and 17/25733-1, respectively. AH received support from DAAD, Cambridge Trust and Emmanuel College, Cambridge. GALZ received a fellowship from the Support Program for Foreign Ph.D. Student and Ibero-American Graduate Association (PAEDEx/AUIP 2014. TK receives an EPSRC Ph.D. fellowship and MVBD receives a CNPq research fellowship (Bolsa de Produtividade em Pesquisa, Level 2). We thank LNLS and LNBio at CNPEM in Campinas for allowing the use of the MX2 and NMR facilities, respectively; C. Oliveira and I. Cuccuvia from Institute of Physics and Institute of Chemistry, University of São Paulo, respectively for allowing the access of calorimetry facilities.

