## Supplementary material for "Using a fragment-based approach to identify novel chemical scaffolds targeting the dihydrofolate reductase (DHFR) from *Mycobacterium tuberculosis*"

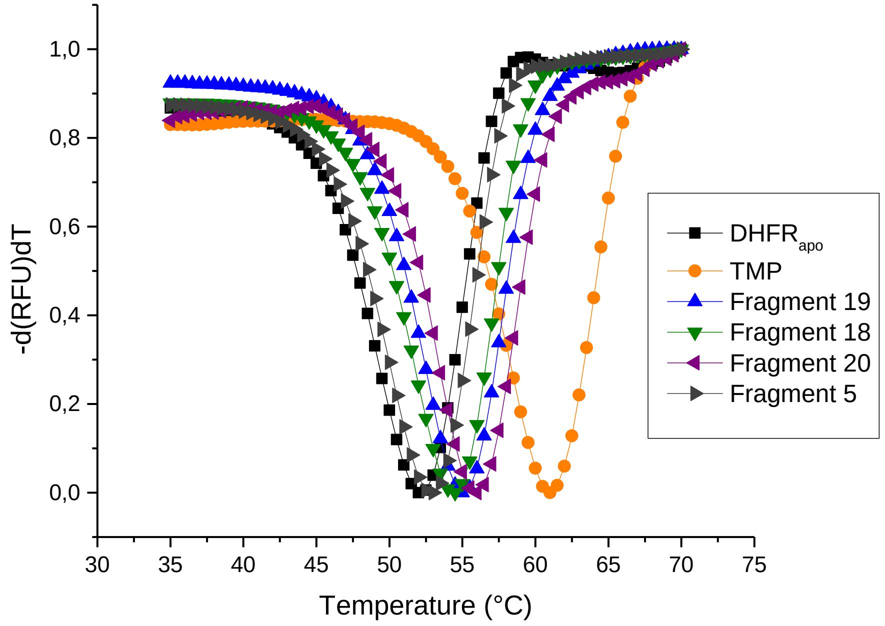


**Supplementary Figure 1.** Thermal shift (DSF) analysis of *Mt*DHFR (3 µM) in the presence of 5% of DMSO and 1mM of NADPH (holoenzyme). Selected fragments and with trimethoprim (TMP) (5mM).

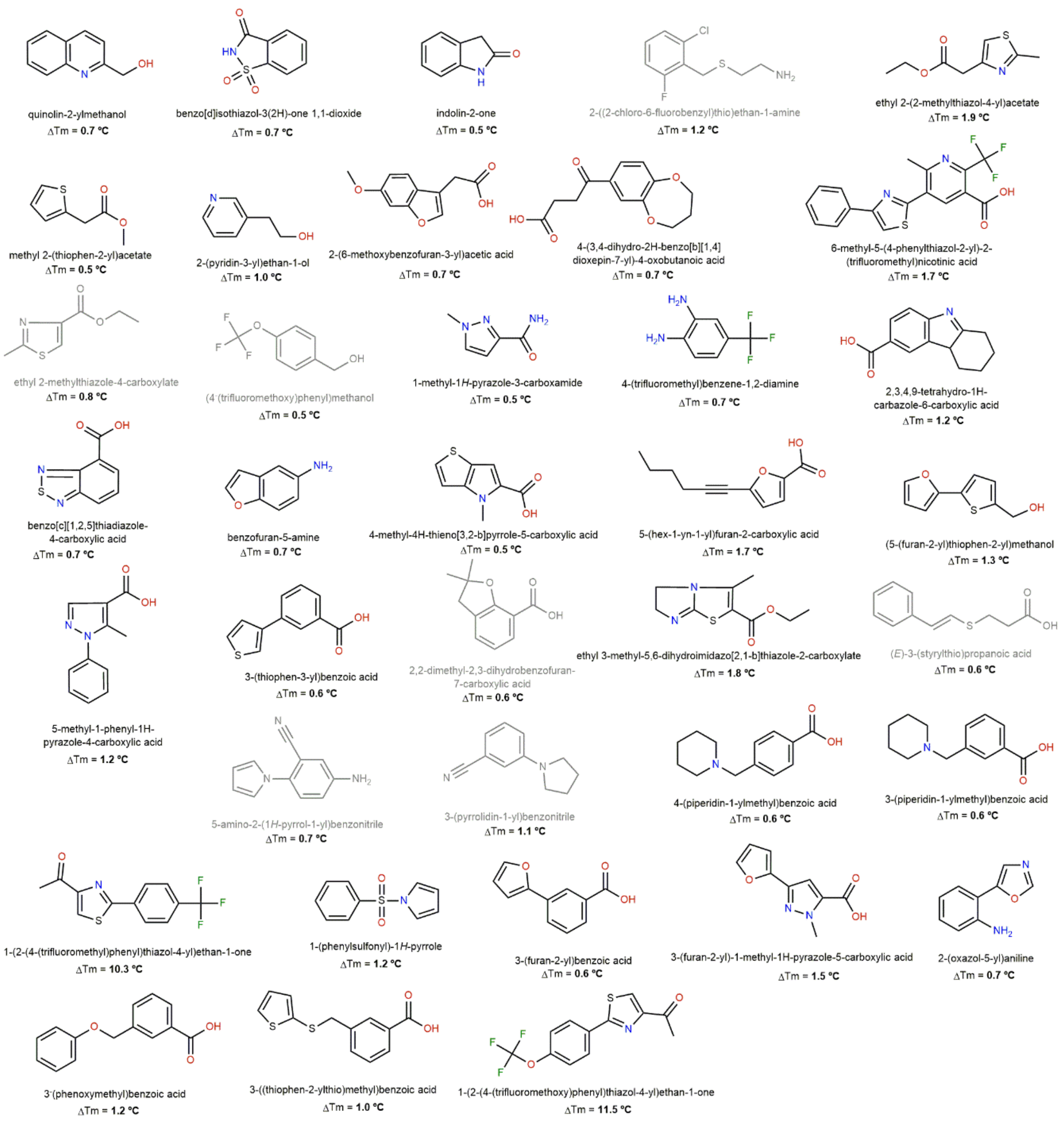

quinolin-2-ylmethanol
ΔTm = 0.7 °C
benzo[d]isothiazol-3(2H)-one 1,1-dioxide
ΔTm = 0.7 °C
indolin-2-one
ΔTm = 0.5 °C
2-((2-chloro-6-fluorobenzyl)thio)ethan-1-amine
ΔTm = 1.2 °C
ethyl 2-(2-methylthiazol-4-yl)acetate
ΔTm = 1.9 °C
methyl 2-(thiophen-2-yl)acetate
ΔTm = 0.5 °C
2-(pyridin-3-yl)ethan-1-ol
ΔTm = 1.0 °C
2-(6-methoxybenzofuran-3-yl)acetic acid
ΔTm = 0.7 °C
4-(3,4-dihydro-2H-benzo[b][1,4]
dioxepin-7-yl)-4-oxobutanoic acid
ΔTm = 0.7 °C
6-methyl-5-(4-phenylthiazol-2-yl)-2-
(trifluoromethyl)nicotinic acid
ΔTm = 1.7 °C
ethyl 2-methylthiazole-4-carboxylate
ΔTm = 0.8 °C
(4-(trifluoromethoxy)phenyl)methanol
ΔTm = 0.5 °C
1-methyl-1H-pyrazole-3-carboxamide
ΔTm = 0.5 °C
4-(trifluoromethyl)benzene-1,2-diamine
ΔTm = 0.7 °C
2,3,4,9-tetrahydro-1H-
carbazole-6-carboxylic acid
ΔTm = 1.2 °C
benzo[c][1,2,5]thiadiazole-
4-carboxylic acid
ΔTm = 0.7 °C
benzofuran-5-amine
ΔTm = 0.7 °C
4-methyl-4H-thieno[3,2-b]pyrrole-5-carboxylic acid
ΔTm = 0.5 °C
5-(hex-1-yn-1-yl)furan-2-carboxylic acid
ΔTm = 1.7 °C
(5-(furan-2-yl)thiophen-2-yl)methanol
ΔTm = 1.3 °C
5-methyl-1-phenyl-1H-
pyrazole-4-carboxylic acid
ΔTm = 1.2 °C
3-(thiophen-3-yl)benzoic acid
ΔTm = 0.6 °C
2,2-dimethyl-2,3-dihydrobenzofuran-
7-carboxylic acid
ΔTm = 0.6 °C
ethyl 3-methyl-5,6-dihydroimidazo[2,1-b]thiazole-2-carboxylate
ΔTm = 1.8 °C
(E)-3-(styrylthio)propanoic acid
ΔTm = 0.6 °C
5-amino-2-(1H-pyrrol-1-yl)benzonitrile
ΔTm = 0.7 °C
3-(pyrrolidin-1-yl)benzonitrile
ΔTm = 1.1 °C
4-(piperidin-1-ylmethyl)benzoic acid
ΔTm = 0.6 °C
3-(piperidin-1-ylmethyl)benzoic acid
ΔTm = 0.6 °C
1-(2-(4-(trifluoromethyl)phenyl)thiazol-4-yl)ethan-1-one
ΔTm = 10.3 °C
1-(phenylsulfonyl)-1H-pyrrole
ΔTm = 1.2 °C
3-(furan-2-yl)benzoic acid
ΔTm = 0.6 °C
3-(furan-2-yl)-1-methyl-1H-pyrazole-5-carboxylic acid
ΔTm = 1.5 °C
2-(oxazol-5-yl)aniline
ΔTm = 0.7 °C
3-(phenoxymethyl)benzoic acid
ΔTm = 1.2 °C
3-((thiophen-2-ylthio)methyl)benzoic acid
ΔTm = 1.0 °C
1-(2-(4-(trifluoromethoxy)phenyl)thiazol-4-yl)ethan-1-one
ΔTm = 11.5 °C

**Supplementary Figure 2.** Fragment-hits identified by Differential Scanning Fluorometry (DSF). In bold is shown the $\Delta T_m$ obtained in comparison to the holoenzyme. In gray are the fragments that were not further tested by STD.

1

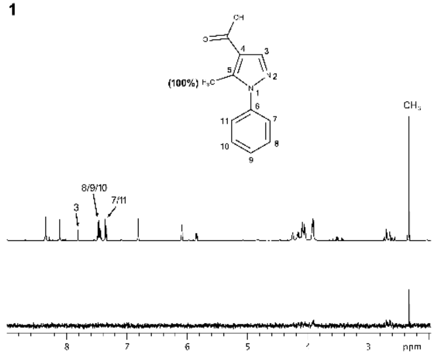

2

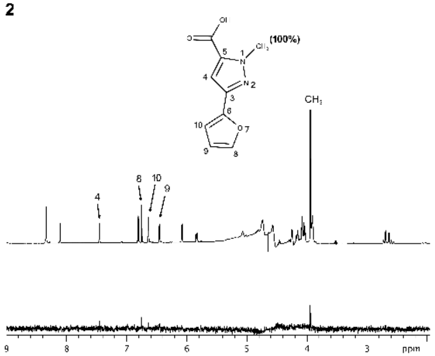

3

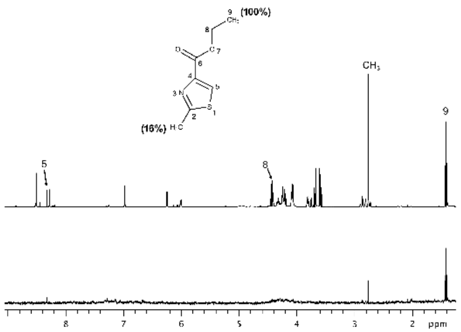

4

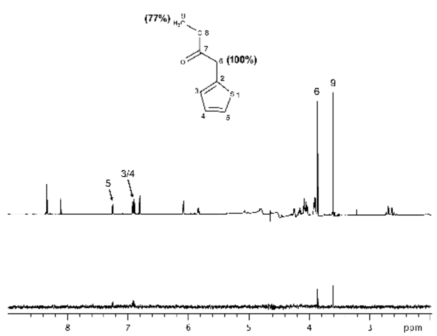

5

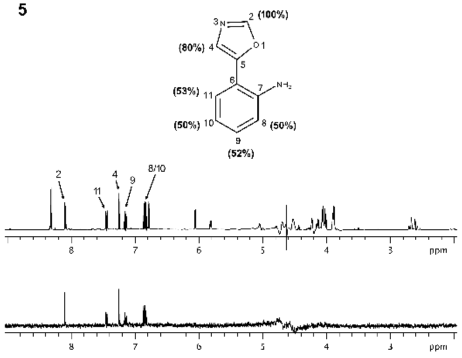

6

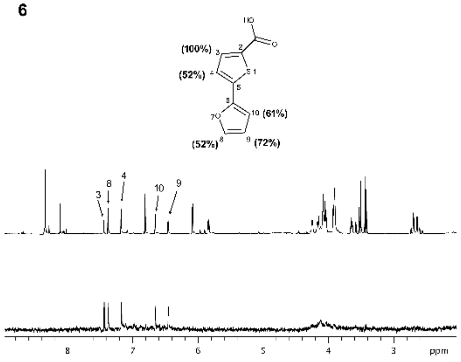

7

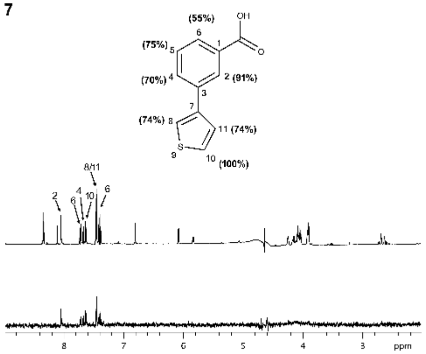

8

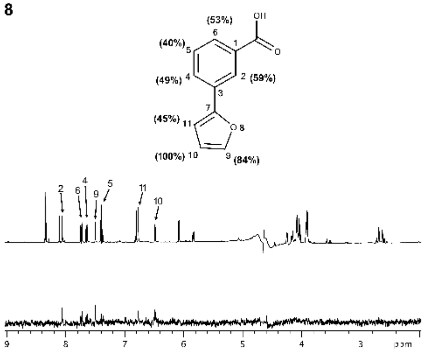

9

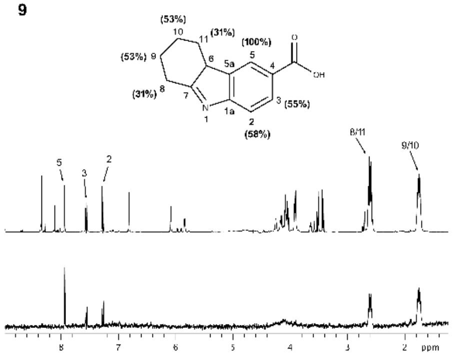

10

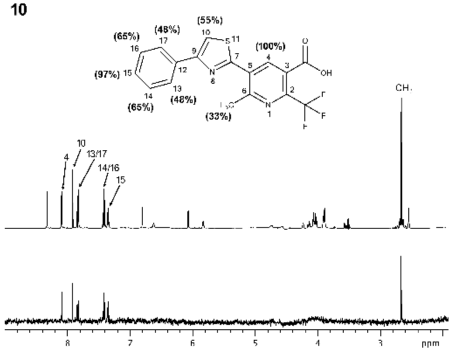

11

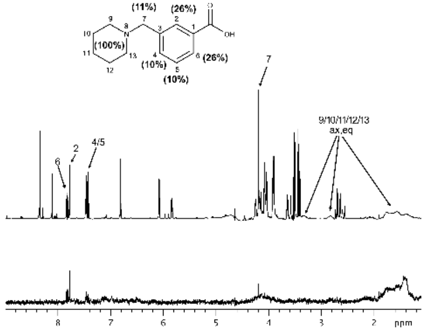

12

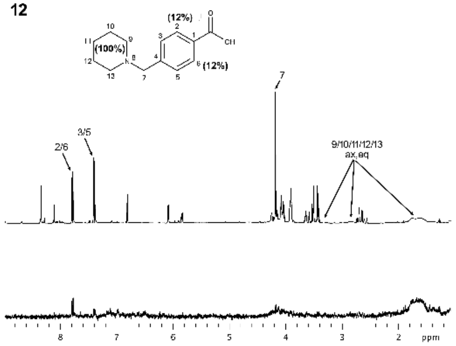

**13**

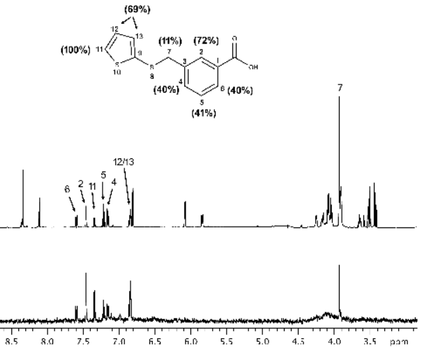


**14**

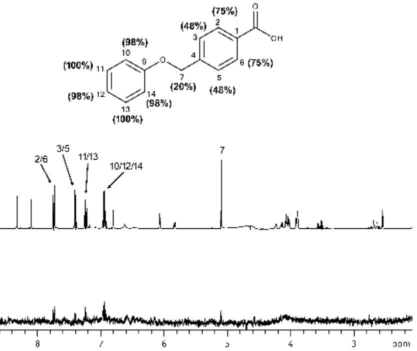


**19**

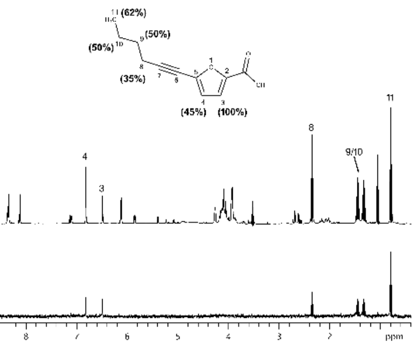


**20**

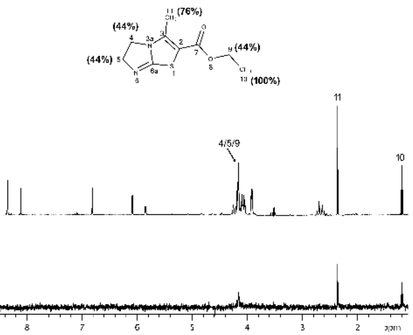


**15**

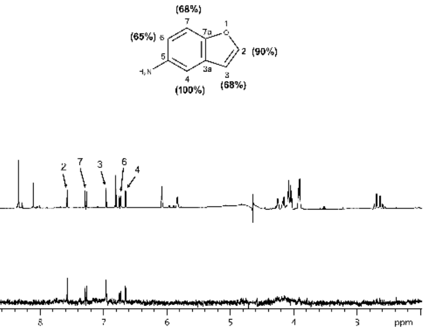


**16**

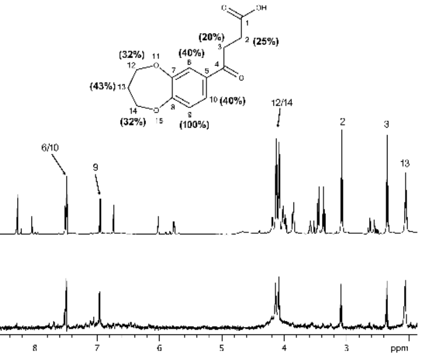


**21**

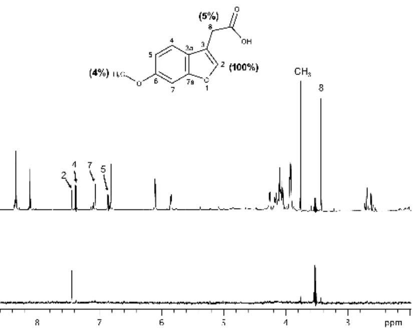


**17**

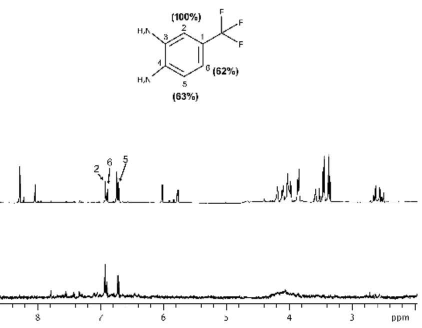


**18**

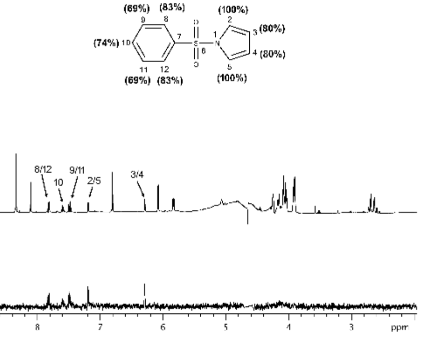

**Supplementary Figure 3.** Saturation Transfer Difference **(**STD)-NMR analysis of the positive fragments

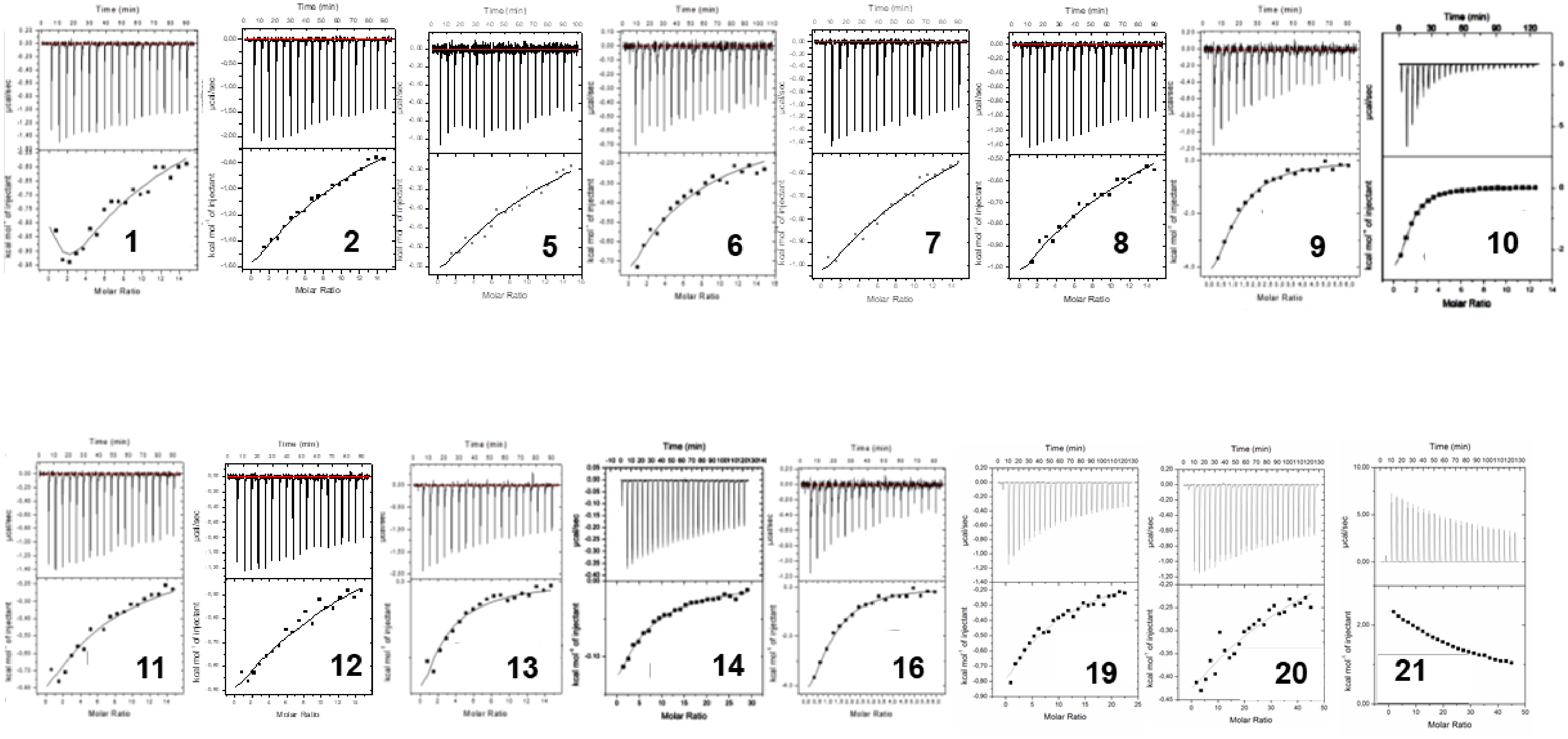


**Supplementary Figure 4.** Isotherms obtained by ITC assays for the interaction of the fragments with MtDHFR

**Table 1.** Calorimetric analysis of fragments against MtDHFR

| Fragment | ΔG (kcal/mol) | ΔH (kcal/mol) | (-) TΔS (kcal/mol) | Kd (mM) | LE |
|---|---|---|---|---|---|
| **1** | -4,28 | -4.69 ± 0.25 | 0,41 | 0.64 ± 0.15 | 0.28 |
| **2** | -0.12 | -9.83±0.45 | 9.71 | 8.6±0.4 | -- |
| **5** | -0.24 | -5.93±1.46 | 5.69 | 14.8±4.2 | -- |
| **6** | -3.67 | -13.87 ± 0.81 | 10.19 | 2.5 ± 0.2 | 0.25 |
| **7** | -0.05 | -10.0±1.28 | 9.95 | 9.5±0.8 | -- |
| **8** | -0.15 | -7.85±1.16 | 7.70 | 9.3±1.4 | -- |
| **9** | -5.45 | -4.72 ± 0.203 | -0.73 | 0.098 ± 0.02 | 0.34 |
| **10** | -5.50 | -3.94 ± 0.154 | -1.53 | 0.095 ± 0.005 | 0.23 |
| **11** | -3.37 | -18.93 ± 1.994 | 15.56 | 3.2 ± 0.4 | 0.21 |
| **12** | -0.33 | -7.78±1.36 | 7.45 | 9.6±1.5 | -- |
| **13** | -4.19 | -8.75 ± 0.586 | 4.57 | 0.75 ± 0.09 | 0.26 |
| **14** | -4.46 | -2.11 ± 0.045 | -2.35 | 0.50 ± 0.02 | 0.26 |
| **16** | -3.45 | -16.11 ± 1.759 | 12.65 | 2.7 ± 0.4 | 0.19 |
| **19** | -0.42 | -2.23±0.94 | 1.81 | 1.1±0.07 | -- |
| **20** | -0.31 | -9.22±1.16 | 8.91 | 8.7±1.3 | -- |
| **21** | -0.33 | 30.66±4.2 | 30.99 | 5.1±0.9 | -- |

**Supplementary Table 2. Data collection and refinement statistics for the MtDHFR in complex with fragments**

| | Frag 1 | Frag 2 | Frag 3 | Frag 10 | Frag 11 | Frag 13 | Frag 14 | Frag 16 | Frag 17 |
|---|---|---|---|---|---|---|---|---|---|
| **PDB entry** | 6VS5 | 6VS6 | 6VS8 | 6VVB | 6VS9 | 6VSD | 6VSE | 6VSF | 6VSG |
| **Wavelength** | 1.458680 | 1.458680 | 1.458680 | 0.97950 | 1.458680 | 1.458680 | 1.458680 | 1.458680 | 1.458680 |
| **Resolution range** | 35.23 - 1.76 (1.82 - 1.76) | 31.87 - 1.85 (1.92 - 1.85) | 30.94 - 1.83 (1.89 - 1.83) | 26.99 - 1.45 (1.50 - 1.45) | 35.29 - 1.84 (1.91 - 1.84) | 46.15 - 2.30 (2.39 - 2.30) | 10.99 - 1.76 (1.82 - 1.76) | 46.48 - 2.01 (2.08 - 2.01) | 46.15 - 2.30 (2.37 - 2.30) |
| **Space group** | P 21 21 21 | P 21 21 21 | P 21 21 21 | C 1 2 1 | P 21 21 21 | P 21 21 21 | P 21 21 21 | P 21 21 21 | P 21 21 21 |
| **Unit cell** | 61.48 70.45 71.88 90 90 90 | 61.16 70.58 71.43 90 90 90 | 60.28 70.78 72.11 90 90 90 | 67.86 73.73 36.78 90 99.89 90 | 61.01 70.58 72.20 90 90 90 | 60.91 70.70 72.13 90 90 90 | 61.44 71.11 72.11 90 90 90 | 61.56 70.90 71.94 90 90 90 | 60.91 70.70 72.13 90 90 90 |
| **Total reflections** | 372719 (14220) | 266847 (9787 | 328044 (14099) | 191519 (27836) | 337371 (14672) | 409192 (12140) | 391509 (15437) | 261901 (17499) | 174536 (14704) |
| **Unique reflections** | 31570 (3008) | 26631 (2380) | 27734 (2609) | 27829 (2555) | 27448 (2602) | 14181 (1272) | 31765 (3000) | 21357 (2055) | 14181 (1272) |
| **Multiplicity** | 11.8 (8.6) | 10.0 (7.4) | 11.8 (9.0) | 6.9 (6.8) | 12.3 (9.5) | 11.4 (7.4) | 12.2 (9.3) | 12.2 (11.5) | 12.3 (11.8) |
| **Completeness (%)** | 98.90 (96.75) | 98.94 (90.06) | 99.36 (95.01) | 88.00 (81.0) | 99.32 (95.14) | 99.15 (91.58) | 98.95 (95.33) | 99.27 (97.44) | 99.15 (91.58) |
| **Mean I/sigma(I)** | 19.0 (2.2) | 12.5 (2.4) | 19.1 (2.2) | 15.3 (3.5) | 17.7 (2.1) | 21.0 (1.8) | 16.1 (2.2) | 11.1 (2.1) | 13.2(2.6) |
| **Wilson B-factor** | 20.77 | 20.98 | 21.35 | 14.91 | 23.67 | 29.60 | 21.10 | 20.34 | 29.60 |
| **R-merge** | 0.08 (0.89) | 0.13 (0.68) | 0.11 (1.1) | 0.06 (0.47) | 0.10 (1.12) | 0.07 (0.91) | 0.1 (0.98) | 0.23 (1.53) | 0.17(0.99) |
| **R-meas** | 0.09 (1.01) | 0.14 (0.78) | 0.12 (1.22) | 0.08 (0.55) | 0.11 (1.26) | 0.07 (1.04) | 0.11 (1.1) | 0.25 (1.68) | 0.19 (1.08) |
| **R-pim** | 0.03 (0.337) | 0.04 (0.26) | 0.04 (0.40) | 0.04 (0.3) | 0.03 (0.39) | 0.02 (0.37) | 0.03 (0.35) | 0.07 (0.49) | 0.05 (0.31) |
| **CC1/2** | 0.999 (0.716 | 0.996 (0.802) | 0.999 (0.654) | 0.998 (0.909) | 0.999 (0.749) | 0.999 (0.690) | 0.998 (0.736) | 0.995 (734) | 0.996 (0.778) |
| **Reflections used in refinement** | 31361 (3008) | 26622 (2375) | 27729 (2608) | 27829 (2555) | 27440 (2602) | 14180 (1272) | 31750 (3000) | 21354 (2056) | 14180 (1272) |
| **Reflections used for R-free** | 1601 (168) | 1273 (98) | 1463 (159) | 1388 (129) | 1392 (126) | 674 (54) | 1615 (169) | 1115 (93) | 674 (54) |
| **R-work** | 0.206 (0.316) | 0.235 (0.318) | 0.170 (0.245) | 0.169 (0.217) | 0.18 (0.257) | 0.175 (0.229) | 0.200 (0.224) | 0.163 (0.210) | 0.175 (0.229) |
| **R-free** | 0.266 (0.355) | 0.272 (0.327) | 0.221 (0.272) | 0.193 (0.257) | 0.208 (0.268) | 0.228 (0.287) | 0.248 (0.266) | 0.211 (0.282) | 0.228 (0.286) |
| **Number of non-hydrogen atoms** | 2948 | 2902 | 2982 | 1506 | 2890 | 2800 | 2785 | 2960 | 2800 |
| **macromolecules** | 2484 | 2492 | 2490 | 1257 | 2488 | 2484 | 2483 | 2488 | 2484 |
| **ligands** | 173 | 136 | 155 | 95 | 156 | 116 | 120 | 179 | 116 |

| | | | | | | | | | |
|---|---|---|---|---|---|---|---|---|---|
| **solvent** | 291 | 274 | 337 | 154 | 246 | 200 | 182 | 293 | 200 |
| **Protein residues** | 318 | 318 | 318 | 161 | 318 | 318 | 318 | 318 | 318 |
| **RMS(bonds)** | 0.008 | 0.008 | 0.008 | 0.005 | 0.007 | 0.008 | 0.007 | 0.009 | 0.008 |
| **RMS(angles)** | 1.12 | 1.06 | 1.07 | 0.84 | 1.11 | 0.94 | 1.10 | 1.12 | 0.94 |
| **Ramachandran favored (%)** | 97.13 | 96.50 | 96.50 | 98.10 | 97.45 | 96.82 | 97.77 | 97.13 | 96.82 |
| **Ramachandran allowed (%)** | 2.87 | 3.18 | 3.18 | 1.90 | 2.23 | 2.87 | 2.23 | 2.87 | 2.87 |
| **Ramachandran outliers (%)** | 0.00 | 0.32 | 0.32 | 0.00 | 0.32 | 0.32 | 0.00 | 0.00 | 0.32 |
| **Rotamer outliers (%)** | 0.79 | 0.39 | 0.78 | 0.00 | 0.39 | 0.00 | 0.79 | 0.39 | 0.00 |
| **Clashscore** | 6.99 | 11.30 | 7.58 | 2.70 | 5.24 | 10.39 | 4.52 | 4.83 | 10.39 |
| **Average B-factor** | 28.45 | 26.36 | 26.99 | 24.19 | 28.54 | 31.37 | 26.29 | 22.77 | 31.37 |
| **macromolecules** | 27.58 | 25.34 | 24.99 | 22.56 | 27.56 | 30.69 | 25.85 | 21.47 | 30.69 |
| **ligands** | 25.61 | 27.04 | 37.82 | 22.69 | 33.70 | 33.50 | 24.48 | 24.96 | 33.50 |
| **solvent** | 37.54 | 35.27 | 36.81 | 38.42 | 35.25 | 38.60 | 33.44 | 32.50 | 38.60 |
| **Number of TLS groups** | 1 | 1 | 1 | 1 | 0 | 1 | 1 | 1 | 1 |

Statistics for the highest-resolution shell are shown in parentheses.

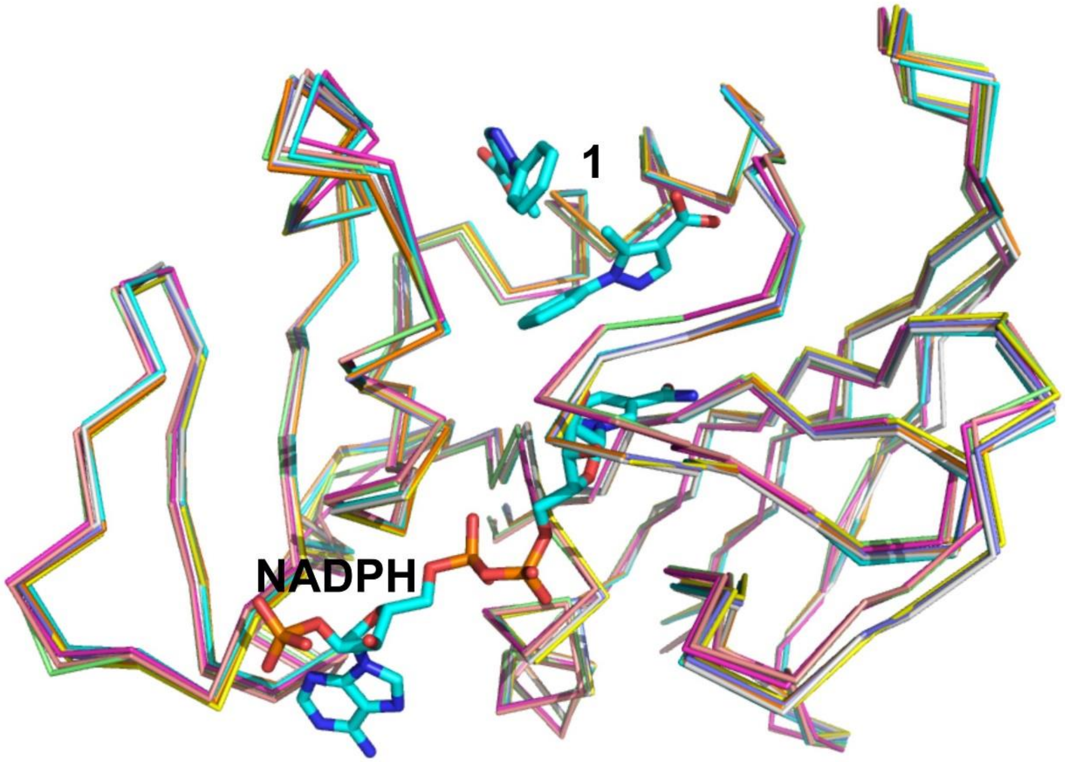


**Supplementary Figure 5.** Superposition of all structures of MtDHFR:NADPH in complex with fragments. The NADPH and the ligand of the complex MtDHFR:NADPH:**1** is shown to identify the cofactor and fragment binding sites, respectively.

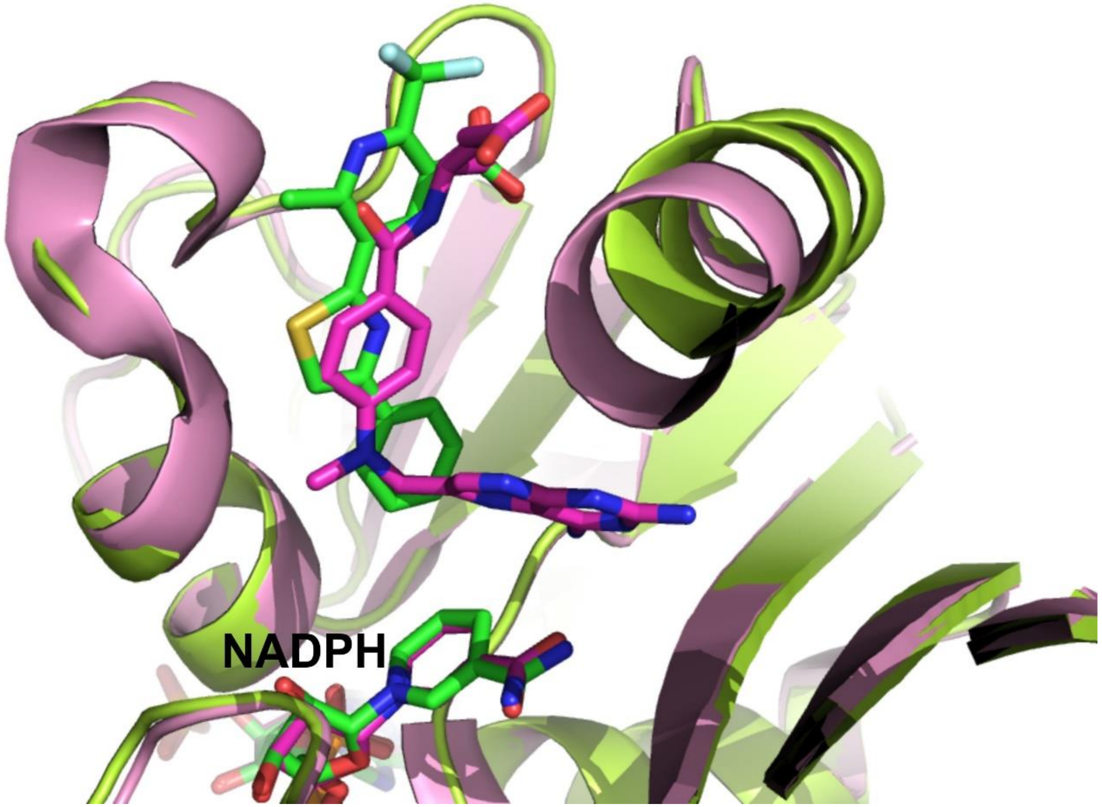


**Supplementary Figure 6.** Superposition of the complexes MtDHFR-NADPH-**10** (in green) and MtDHFR-NADPH-MTX (in purple).

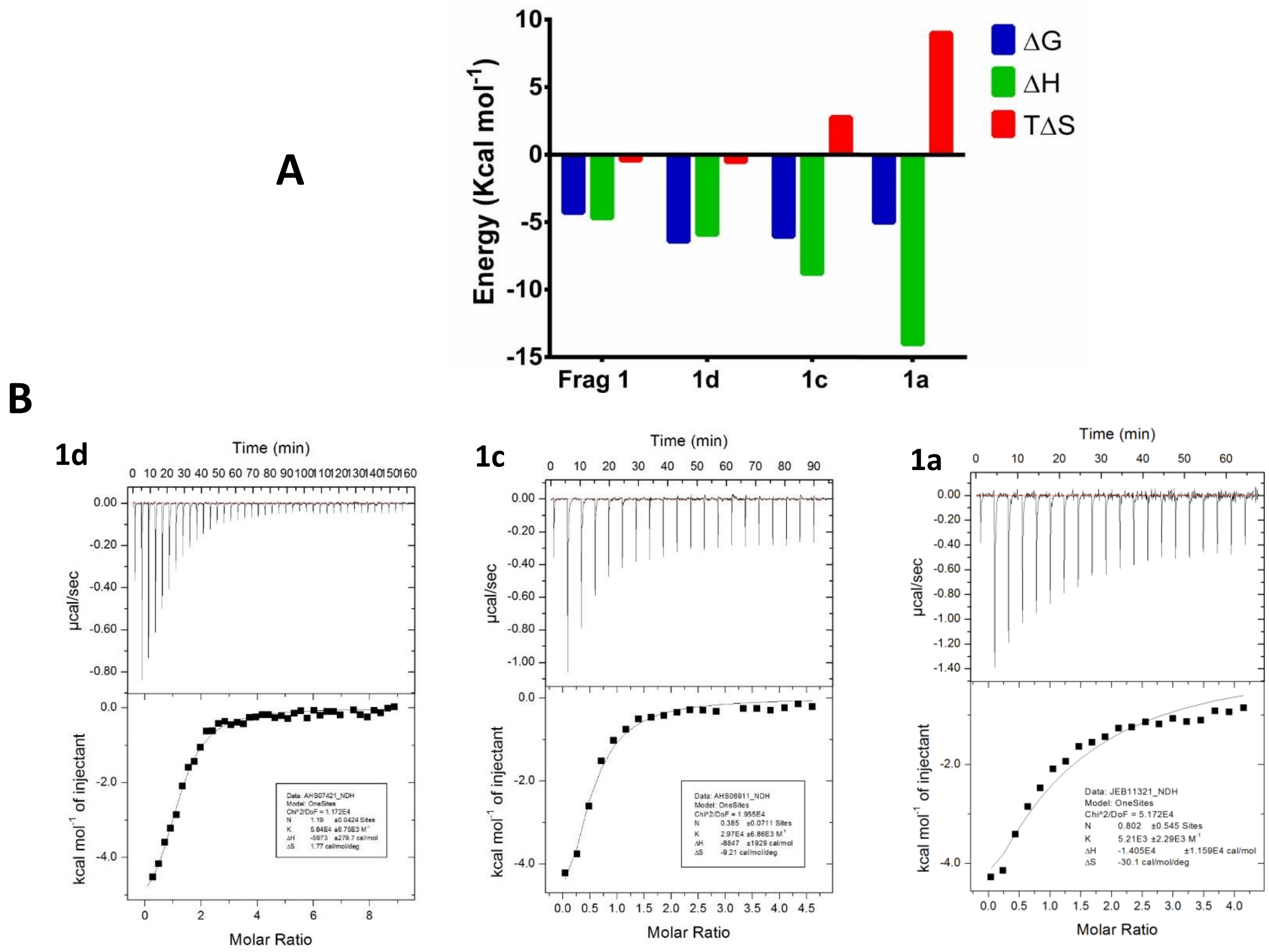


S**upplementary figure 7**. Thermodynamic analysis for the interaction of MtDHFR:NADPH with compounds based on fragment 1. (A) Histogram showing the enthalpic and entropic components of the interaction of different compounds (B) Thermograms obtained for three of the compounds based on Fragment **1**.

**Supplementary Table 3. Data collection and refinement statistics for the MtDHFR in complex with compounds based on fragment 1.**

| | **1a** | **1b** | **1c** | **1d** |
|---|---|---|---|---|
| **PDB entry** | 6VV6 | 6VV9 | 6VV7 | 6VV8 |
| **Wavelength** | 1.458690 | 1.458690 | 1.458690 | 1.458690 |
| **Resolution range** | 11.43 - 2.05 (2.12 - 2.05) | 31.8 - 2.18 (2.26 - 2.18) | 12.73 - 2.0 (2.07 - 2.0) | 35.75 - 2.68 (2.78 - 2.68) |
| **Space group** | P 21 21 21 | P 21 21 21 | P 21 21 21 | P 21 21 21 |
| **Unit cell** | 61.49 71.66 72.20 90 90 90 | 61.18 70.87 72.08 90 90 90 | 61.50 72.02 72.04 90 90 90 | 61.17 71.49 72.21 90 90 90 |
| **Total reflections** | 170979 (13433) | 152267 (13505 | 285475 (40885) | 115065 (14140) |
| **Unique reflections** | 20447 (2008) | 16880 (1647) | 22136 (2162) | 9274 (898) |
| **Multiplicity** | 8.3 (8.6) | 9.0 (9.4) | 12.8 (12.8) | 12.3 (12.0) |
| **Completeness (%)** | 99.22 (100.00) | 99.75 (100.00) | 99.42 (99.22) | 99.83 (99.23) |
| **Mean I/sigma(I)** | 10.9 (2.1) | 11.0 (2.0) | 13.9 (3.0) | 10.8 (2.8) |
| **Wilson B-factor** | 33.09 | 36.40 | 29.60 | 37.51 |
| **R-merge** | 0.12 (0.98) | 0.13 (1.07 | 0.13 (0.82) | 0.24 (1.07) |
| **R-meas** | 0.14 (1.1) | 0.15 (1.19 | 0.13 (0.86) | 0.26 (1.16) |
| **R-pim** | 0.06 (0.51) | 0.05 (0.52) | 0.04 (0.24) | 0.1 (0.46) |
| **CC1/2** | 0.997 (0.751) | 0.997 (0.713) | 0.997 (0.759) | 0.993 (0.797) |
| **Reflections used in refinement** | 20446 (2008) | 16865 (1648) | 22136 (2162) | 9271 (897) |
| **Reflections used for R-free** | 1065 (101) | 822 (82) | 1048 (86) | 423 (52) |
| **R-work** | 0.236 (0.313) | 0.245 (0.321) | 0.314 (0.367) | 0.236 (0.282) |
| **R-free** | 0.263 (0.284) | 0.281 (0.357) | 0.331 (0.402) | 0.293 (0.427) |
| **Number of non-hydrogen atoms** | 2750 | 2740 | 2738 | 2681 |
| **macromolecules** | 2488 | 2488 | 2488 | 2488 |
| **ligands** | 169 | 179 | 179 | 154 |
| **solvent** | 93 | 73 | 71 | 39 |
| **Protein residues** | 320 | 320 | 318 | 318 |
| **RMS(bonds)** | 0.019 | 0.019 | 0.019 | 0.010 |
| **RMS(angles)** | 1.40 | 1.38 | 1.42 | 1.07 |
| **Ramachandran favored (%)** | 95.86 | 97.45 | 98.09 | 93.31 |
| **Ramachandran allowed (%)** | 4.14 | 2.55 | 1.91 | 6.69 |
| **Ramachandran outliers (%)** | 0.00 | 0.00 | 0.00 | 0.00 |
| **Rotamer outliers** | 0.00 | 0.00 | 0.00 | 2.36 |

| | | | | |
|---|---|---|---|---|
| **(%)** | | | | |
| **Clashscore** | 4.27 | 5.23 | 5.61 | 14.97 |
| **Average B-factor** | 36.38 | 42.12 | 55.42 | 32.84 |
| | 35.95 | 40.81 | 57.58 | 32.73 |
| **macromolecules** | | | | |
| **ligands** | 43.40 | 62.01 | 32.71 | 35.97 |
| **solvent** | 35.38 | 37.89 | 36.78 | 27.93 |
| **Number of TLS groups** | 1 | 1 | 1 | |

Statistics for the highest-resolution shell are shown in parentheses.

### Supplementary Experimental Section

#### *Differential Scanning Fluorimetry against a library based on fragment **1***

The Abell research group has an in-house library of compounds and the scaffolds were selected based on the fragment hit **1**. About 300 molecules based on fragment **1** screened by DSF. All compounds screened had a purity of >95% by LCMS analysis. The thermal shift assay was performed in 96-well plates. Each well-contained DHFR (3 μM), 1 mM NADPH, 5 X SYPRO Orange and fragments (1 mM or 5 mM) in 20 mM phosphate buffer, containing 50 mM KCl, pH 7.0. The total sample volume was 50 μL with a final DMSO-$d_6$ concentration of 5% v/v. Plates were centrifuged at 1000 rpm for 30 seconds. Differential scanning fluorimetry was conducted on a CFX Connect™ instrument (BioRad). The temperature was increased from 25 – 95 °C at a rate of 0.5 °C/min. The data were processed using Microsoft Excel. The fluorescence intensity was plotted versus the temperature. The melting point ($T_m$), defined as the inflection point, was identified from the minima of each curve's first derivative. The thermal shift ($\Delta T_m$) in the presence of fragments was calculated relative to DMSO control wells.

#### *Isothermal Titration Calorimetry of compounds identified based on **1***

ITC experiments were performed on a MicroCal auto-iTC200 (Malvern Instruments, Malvern UK) instrument at 25 °C. Ligand solutions (0.5 – 4 mM) and protein solutions (50 – 120 μM) were prepared in the same buffer (20 mM phosphate, 50 mM KCl, pH = 7.0, containing 1 mM NADPH). The final DMSO-$d_6$ concentration was varied between 0.5 – 5% v/v, depending on the solubility of the ligand. Titrations were typically 19 × 2.0 μL injections with 200 – 250s spacing time. An initial 0.2 μL injection was discarded during data analysis. Background heats measured by a control titration of ligand solution into a buffer without protein were subtracted from the raw data. Titration data were analysed and fitted to a one-site binding model using Origin analysis software provided by the manufacturer. The stoichiometry was set to N = 1 or allowed to vary when high goodness of fit could be obtained.
